# A quantitative view of strategies to engineer cell-selective ligand binding

**DOI:** 10.1101/2020.11.25.398974

**Authors:** Zhixin Cyrillus Tan, Brian Orcutt-Jahns, Aaron S. Meyer

**Affiliations:** Bioinformatics Interdepartmental Program, University of California, Los Angeles; Department of Bioengineering, University of California, Los Angeles; Jonsson Comprehensive Cancer Center, University of California, Los Angeles; Eli and Edythe Broad Center of Regenerative Medicine and Stem Cell Research, University of California, Los Angeles

## Abstract

A critical property of many therapies is their selective binding to specific target populations. Exceptional specificity can arise from high-affinity binding to unique cell surface targets. In many cases, however, therapeutic targets are only expressed at subtly different levels relative to off-target cells. More complex binding strategies have been developed to overcome this limitation, including multi-specific and multi-valent molecules, but these create a combinatorial explosion of design possibilities. Therefore, guiding strategies for developing cell-specific binding are critical to employ these tools. Here, we extend a multi-valent binding model to multi-ligand and multi-receptor interactions. Using this model, we explore a series of mechanisms to engineer cell selectivity, including mixtures of molecules, affinity adjustments, and valency changes. Each of these strategies maximizes selectivity in distinct cases, leading to synergistic improvements when used in combination. Finally, we identify situations in which selectivity cannot be derived through passive binding alone to highlight areas in need of new developments. In total, this work uses a quantitative model to unify a comprehensive set of design guidelines for engineering cell-specific therapies.

**Summary points:**

- Affinity, valency, and other alterations to target cell binding provide enhanced selectivity in specific situations.
- Evidence for the effectiveness and limitations of each strategy are abundant within the drug development literature.
- Combining strategies can offer enhanced selectivity.
- A simple, multivalent ligand-receptor binding model can help to direct therapeutic engineering.

## Introduction

Many drugs both derive their therapeutic benefit and avoid toxicity through selective binding to specific cells within the body. Often, target cells can differ from off-target populations only subtly in surface receptor expression, making selective activation of target cells difficult to achieve. Even with a drug of very specific molecular binding, genetic and non-genetic heterogeneity can create a wide distribution of cell responses. This can result in reduced effectiveness and increased toxicity. Specific cell targeting is a universal challenge in protein-based therapies. For example, in cancer, resistance to anti-tumor antibodies,^1^ targeted inhibitors,^2^ chemotherapies,^3^ and chimeric antigen receptor T cells^4,5^ all can arise through the selection of target cells among heterogeneous cell populations. While the immune system takes advantage of heterogeneity at the single-cell level to translate noisy inflammatory signals into robust yet sensitive responses,^6^ this heterogeneity impedes our effort of creating a highly specific drug. The intricacies of both inter-population receptor expression differences and intra-population receptor expression heterogeneity present significant challenges that limit the selectivity of therapies within the body.

Further improving cell-specific targeting will require new strategies of engineering specificity. Non-cellular therapies such as protein therapies have most extensively been engineered to target specific cell types through mutations that provide high-affinity binding to unique surface antigens.^7^ This strategy can enhance specificity, but only to a limited degree, particularly when target cells can only be distinguished by subtle quantitative differences in surface antigen or by combinations of markers. The limitations of single-antigen targeting have led to efforts to program complex logic into cellular therapies to recognize target cells more specifically.^8–10^ However, non-cellular therapies have considerable benefits in drug access, manufacturing, and reliability; some of the same benefits have begun to be engineered into these agents.

The enormous number of potential configurations of ligand designs make computational tools essential for designing highly selective therapies.^11^ Here, we analyze a suite of molecular approaches for engineering cell-specific binding using a multi-valent, multi-receptor, multi-ligand binding model. We show that strategies including affinity, valency, binding competition, ligand mixtures, and hetero-valent complexes provide distinct improvements in cell-specific targeting. Finally, we combine these strategies to target cells through combinatorial methods. In total, our results demonstrate that multi-valent ligands can offer offer effective cell specific targeting without the need to engineer complex cellular therapies, and that their design can be guided using a mechanistic binding model.

## Results

### A Model System to Explore the Factors Contributing to Cell Selectivity

Here we investigate cell-specific targeting quantitatively by extending a multi-valent, multi-ligand equilibrium binding model. Virtually any therapy, including monoclonal antibodies, small-molecule inhibitors, and cytokines, can be thought of as ligands for respective receptors expressed on target cells. Ligand binding to target cells is a necessary first step, and essential for a drug’s intended action. In contrast, binding to off-target cells can result in unintended effects or toxicity. Some cell populations can be distinguished by their expression of a unique receptor not expressed by other populations, but more commonly, target and off-target cells express the same collection of receptors and differ only in their magnitudes of receptor expression. In such situations, engineering drugs to optimize target cell binding while minimizing their binding to off-target cells is an area of ongoing research, and has inspired a myriad of drug design strategies.^12–14^ In this work, we define cell population selectivity as the ratio of the number of ligands bound to target cell populations divided by the number of ligands bound to off-target cell populations. We will use a quantitative binding estimation for each cell population to examine these strategies.

While ligand-receptor binding events in biology are diverse, they are governed by thermodynamic properties and the law of mass action. For the binding between monomer ligands and receptors, their affinity can be described by the association constant *K*_*a*_, or its reciprocal, the dissociation constant *K*_*d*_. Binding behavior is more complicated when the ligands are multivalent complexes consisting of multiple units, each of which can bind to a receptor (Fig. 1a). During initial association, we assume that the first subunit on a ligand complex binds according to the same dynamics that govern monovalent binding. Subsequent binding events exhibit different behavior, however, due to the increased local concentration of the complex and steric effects. Here, we assume that the effective association constant for the subsequent bindings is proportional to that of the free binding, but scaled by a crosslinking constant, 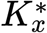, which describes how easily a multivalent ligand bound to a cell monovalently can attain secondary binding. Another consideration that must be made when modeling multivalent binding processes is whether ligand complexes are formed via random assortment of monomers, or whether the monomer composition is uniform across complexes by engineering. We developed a multivalent binding model that calculates the amount of ligand bound at equilibrium taking each of these factors into account (see methods).

**Figure 1:**
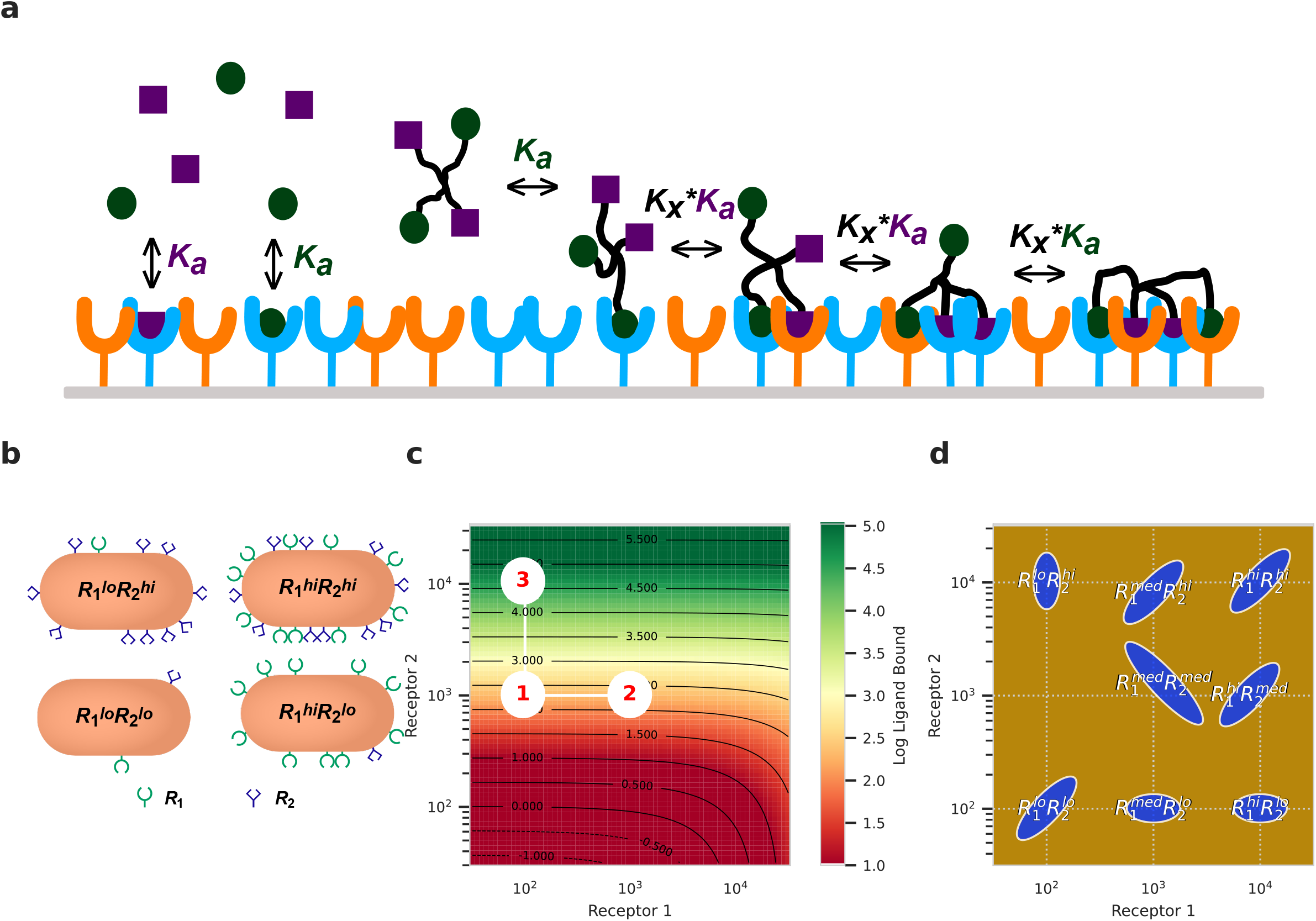
A model system for exploring the factors contributing to cell selectivity. a) A simplified schematic of the binding model. There are two types of receptors and two types of ligand monomers that form a tetravalent complex. b) A cartoon for four cell populations expressing two different receptors at low or high amounts. c) A sample heat/contour map for the model-predicted log ligand bound given the expression of two types of receptors. d) Eight arbitrary theoretical cell populations with various receptor expression profiles.

As a simplification, we will consider theoretical cell populations that express only two receptors capable of binding ligand (Fig. 1b), ranging in abundance from 100 to 10,000 per cell. Figure 1c shows the log-scaled predicted amount of binding of a monovalent ligand given the abundance of two receptors, with the concentration of ligand, *L*_0_, to be 1 nM, and its dissociation constants to the two receptors to be 10 *μ*M and 100 nM, respectively. Because all axes are log-scaled, the number of contour lines between two points indicates the ratio of ligand binding between populations. For instance, in figure 1c, cell populations at points 1 and 2 are on the same contour line and thus have the same amount of ligand bound; the cell populations at points 1 and 3 are separated by multiple contour lines, indicating that cells at point 3 bind more ligands (In fact, the ratio can be read as the exponent of the contour line difference. For point 3 to point 1, the ratio is *e*^4.6−2.3^ ≈ 7.4). Alternatively, we can think of moving from one point to another as a change of expression profile for a cell population. This situation might correspond to some cues inducing expression of a receptor, such as interferon-induced upregulation of MHC and regulatory T cell upregulation of IL-2Rα.^15^ When the amount of receptor 1 (*R*_1_) increases on a cell (a point moves rightward, e.g. from 1 to 2), the amount of binding doesn’t increase significantly. On the contrary, a cell with increased expression of *R*_2_ (moving upward, e.g. from 1 to 3) will bind significantly more ligands. Here, the ligand’s high affinity for *R*_2_ and relatively low affinity for *R*_1_ lead to binding varying more strongly with changes to *R*_2_ expression than *R*_1_.

To analyze more general cases, we arbitrarily defined eight theoretical cell populations according to their differential expression of two receptor types (*R*_1_ and *R*_2_ plotted on x and y axes). As shown in Fig 1d, they either have high (10^4^), medium (10^3^), or low (10^2^) expression of *R*_1_ and *R*_2_. The receptor expression profile within each cell population can also vary widely. To demonstrate cell-to-cell heterogeneity, we also defined intrapopulation variability of each population arbitrarily. For instance, the expression profile of 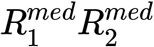 has a wider range. The intrapopulation variance will be accounted for using sigma point filters^16^. We will use this binding model to examine how engineering a ligand using various strategies can improve cell-specific targeting. Although we will only consider two receptor and ligand subunit types respectively, the principles we present can generalize to more complex cases.

### Affinity Provides Selectivity Toward Cell Populations with Divergent Receptor Expression

We first explored altering the ligand binding affinity as an engineering strategy to enhance its cell population specificity. Here, we showed the binding pattern of monovalent ligands with various affinities ranging from 10 *μ*M to 100 nM to both receptors *R*_1_ and *R*_2_ (Fig. 2a). We found that when a target cell population expresses a receptor not expressed by off-target cell populations, enhancing the affinity of the drug to this receptor is a clear and effective strategy to increase selective binding to this population. For example, 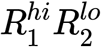 only significantly expresses *R*_1_, while 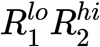 only significantly expresses *R*_2_. When the affinity to *R*_1_ is enhanced and the affinity to *R*_2_ is reduced, the binding selectivity towards 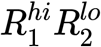 increases (Fig. 2b). The contour plots in Fig. 2a shows this trend more intuitively: when ligand affinity to *R*_1_ increases, which corresponds to shifting from subplots on the left to the ones on the right, binding is shown to vary more strongly according to *R*_1_ expression, indicating an increase in the amount of ligand bound for the populations with high *R*_1_ expression, such as 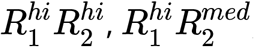, and 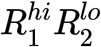.

**Figure 2:**
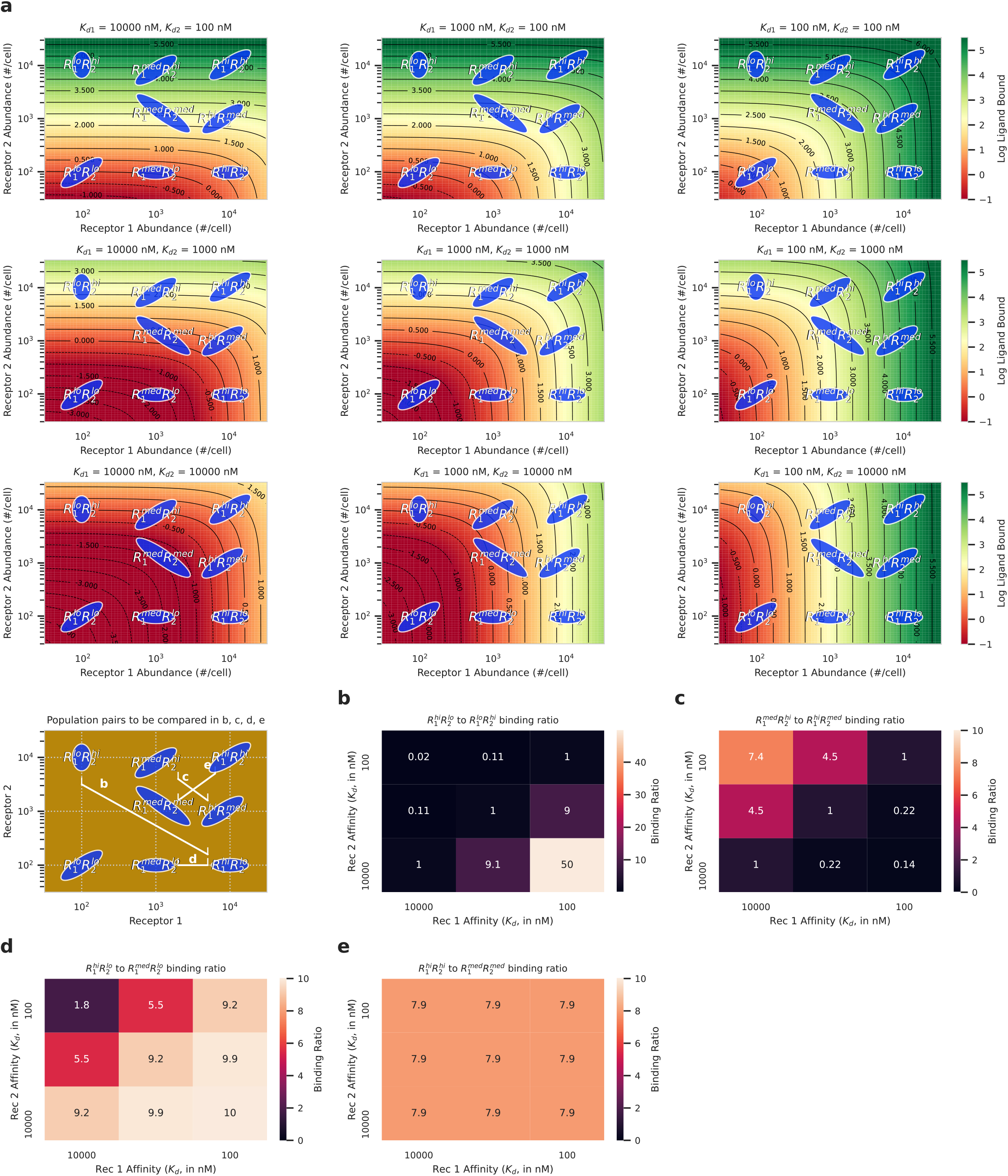
Affinity provides selectivity to cell populations with divergent receptor expression. a) Heat/contour maps of monovalent ligand binding to cell populations given the surface abundance of two receptors. Ligand dissociation constants to these receptors range from 10*μ*M ∼ 100nM. Ligand concentration *L*_0_ = 1nM. b-e) Heatmap of binding ratio of cell populations exposed to a monovalent ligand with dissociation constants to receptor 1 and 2 ranging from 10^4^ ∼ 10^2^nM, at a concentration *L*_0_ = 1nM. Ligand bound ratio of 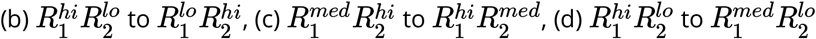 to, and 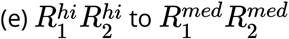.

However, situations where both on- and off-target cell populations express the same set of receptors and differ only in their magnitude of expression are just as common. In these cases, we found out that it is beneficial for the drug to bind tightly to the most comparatively highly expressed receptor on the target population. Some examples of this pattern are the selective binding to 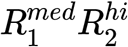 over 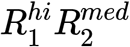, and 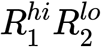 over 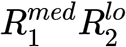 (Fig. 2a). For these two pairs, the benefit of affinity changes is limited by the relative discrepancy in receptor abundances (Fig. 2c,d). Since the ratios of receptor expressions between 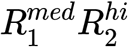 and 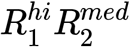 are significantly lower than those between 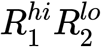 and 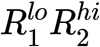, the greatest binding selectivity that can be achieved for these two populations is also significantly lower. When both receptors are uniformally more abundant in a target population than the off target population, such as when comparing 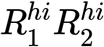 to 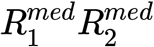, affinity tuning fails to modulate binding selectivity (Fig. 2d). Therefore, to use affinity changes for selectivity enhancement, it is critical to identify which receptors a cell population of interest expresses the most uniquely as compared to off-target populations.

Our binding contour plots are also useful for considering the interplay between affinity modulation and intrapopulation receptor expression heterogeneity. For example, 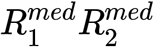 has relatively high variance in both *R*_1_ and *R*_2_ expression (Fig. 2a). When the binding affinities to *R*_1_ and *R*_2_ are divergent, as shown in the subplots in the upper left corner and bottom right corner of Fig. 2a, the ellipse representing its range of receptor abundances rides through multiple contour lines, indicating wide variation in the quantity of bound ligand. This intrapopulation variation in the amount of ligand bound, however, is not observed when the affinities to *R*_1_ and *R*_2_ are more balanced. Thus, a population’s receptor expression heterogeneity and variation are important factors to consider when modulating ligand affinity.

### Valency Enables Selectivity Based on Quantitative Differences in Receptor Abundance

Given the limitations of deriving selectivity from affinity changes, we next explored the effect of valency changes (Fig. 3). Our previous work has shown that this binding model can accurately predict the multivalent binding between IgG antibodies and Fcγ receptors.^17^ To further demonstrate our model’s capacity to predict multivalent binding activity, we fit our model to a set of published experimental measurements wherein fluorescently labeled nanorings were assembled with specific numbers of binding units.^18^ After fitting to determine the crosslinking coefficient of multivalent nanorings 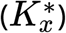 and receptor affinity values (*K*_*a*_), our model was able to accurately match the binding of nanorings with four or eight subunits to cells expressing a known abundance of receptor partners (Fig. S1a).

**Figure 3:**
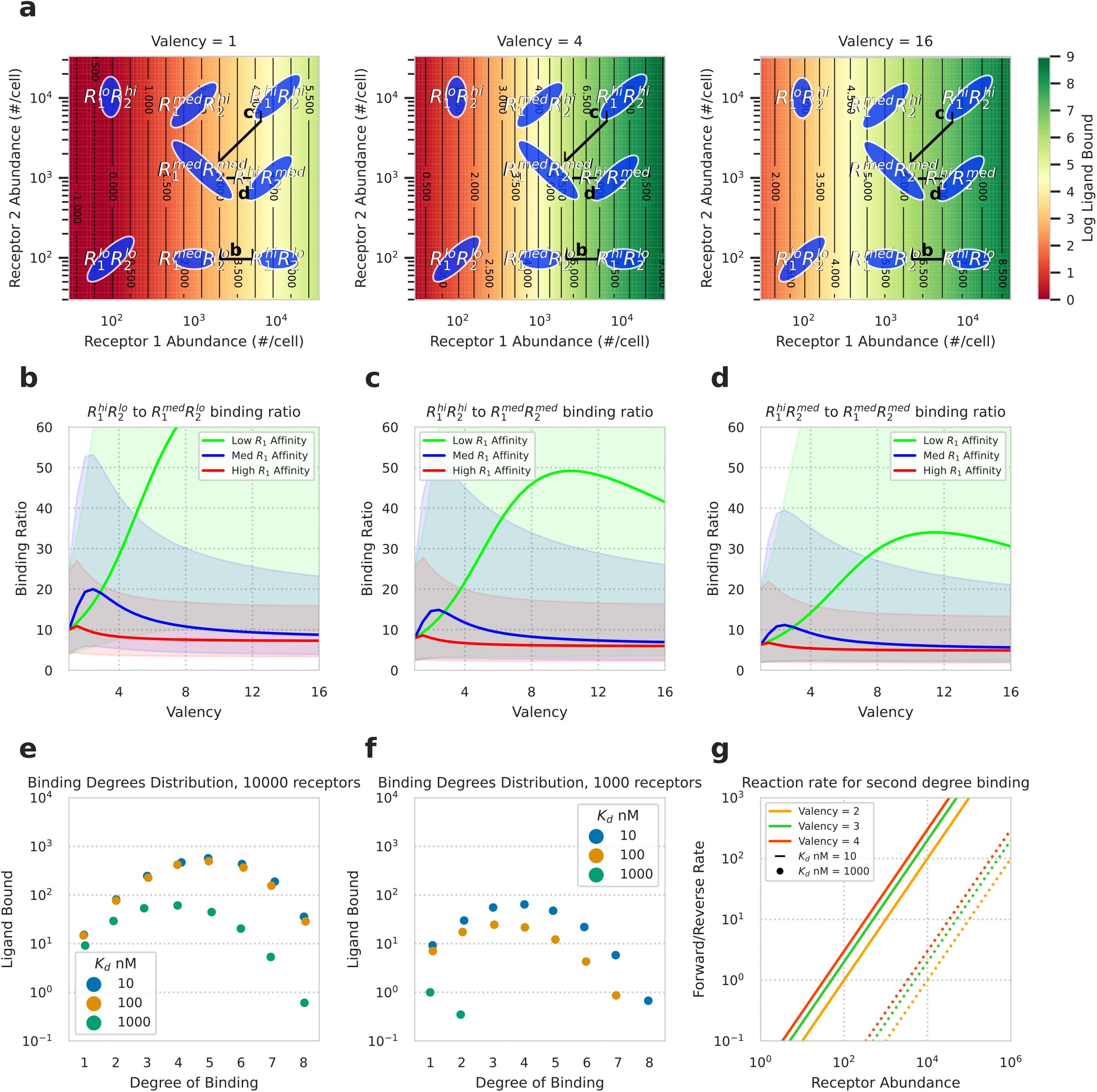
Valency provides selectivity based on receptor expression levels. a) Heat/contour maps of multivalent ligands bound to cell populations given their expression profiles of two receptors. Multivalent ligand subunits bind to only *R*_1_ with an dissociation constant of 100 nM, and do not bind to *R*_2_. Complexes vary in valency from 1 to 16. Ligand concentration *L*_0_ = 1 nM; crosslinking constant 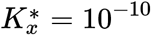. b-d) Ligand binding ratio between various cell populations for ligands of valency ranging from 1 to 16. The shaded areas indicate the variance of binding ratios caused by the intrapopulation heterogeneity and estimated by sigma point filters. b) Ligand bound ratio of 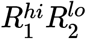 to 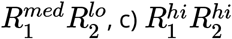 to 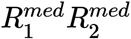, and 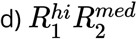 to 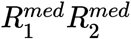. e-f) Number of ligand bound to each possible number of receptors for cells exposed to octavalent ligand complexes composed of subunits with dissociation constants of 1000, 100, or 10 nM for receptor 1. e) Number of octavalent complex bound at each degree for a cell with 10^4^ receptors. f) Number of octavalent complex bound at each degree for a cell with 10^3^ receptors. g) Ratio of forward to reverse binding rate for secondary binding events for multivalent ligands to cells expressing variable amounts of receptors.

Multivalent ligand binding differs from monovalent binding in its nonlinear relationship with receptor density, allowing multivalent ligands to potentially selectively target cells based on receptor abundance.^19^ The effects of varying ligand valency are visualized in Fig. 3a. Here, the ligand only binds to receptor *R*_1_, not receptor *R*_2_, and the valency of the ligand is varied across subplots. Varying ligand valency results in a nonlinear relationship between receptor abundance and binding magnitude. For example, in the monovalent case, there are roughly the same amount of contour lines between 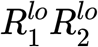 and 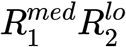 as there are between 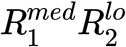 and 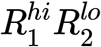. However, in the tetravalent case, there are comparatively more contour lines between 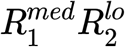 and 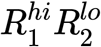, indicating that ligand bound is a nonlinear function of receptor expression when considering multivalent binding (Fig. 3a).

Selectivity derived by changing valency requires coordinate changes in affinity. As many previous studies have suggested, multi-valent ligands can demonstrate particular selectivity to target populations with high receptor abundances when their subunits have low affinity to the receptors.^13,18^ Comparing binding between cell populations exposed to ligands of variable affinities over a range of ligand valency demonstrates the interplay between these factors (Fig. 3b-d). Here, the ligand binding ratios between cell populations are shown for ligands of high, medium, and low affinities for *R*_1_ (*K*_*d*_ of 1000, 100, and 10 nM), and the shaded areas indicate the variance caused by intrapopulation heterogeneity, determined by sigma point filters. The ligand binding ratio between 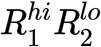 and 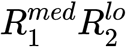 is maximized by low affinity ligands, but requires greater valency to achieve peak binding selectivity when compared to ligands of greater affinity (Fig. 3b). A similar valency optimum for a given affinity is seen for binding selectivity between 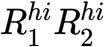 and 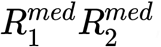, and 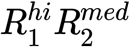 and 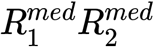 (Fig. 3c-d). Ligands with lower affinities achieve optimal binding with higher valencies and exhibit higher selectivity for cells expressing greater amounts of receptor.

For any specific multivalent ligand at equilibrium, it may bind to any number of receptors up to its valency, which we will refer to as its binding degree. Our model allowed us to identify the mechanism of valency-mediated receptor abundance selectivity by examining the distribution of ligands bound at each degree to different cells for octavalent ligands (valency = 8). A cell expressing 10^4^ receptors displays similar amounts of binding at each degree for ligands with dissociation constants of 1000, 100, and 10 nM (Fig. 3e). However, a cell expressing 10% as many receptors exhibits extremely low amounts of higher-degree binding for ligand complexes of low binding affinity (Fig. 3f). This effect arises due to the ligand’s rate of dissociation being larger than its multivalent binding association rate at low receptor abundances (Fig. 3g). Multivalent ligands undergo initial binding events at rates unaffected by steric effects and changes in local concentration, but low affinity and receptor density severely limit the possibility of secondary binding events. In contrast, cells with a higher abundance of receptors accumulate ligand bound by having them bind multivalently, as the forward rate of secondary binding events is greater than that of receptor-ligand disassociation (Fig. 3f). This effect allows multivalent ligands to achieve selective binding to cells based on their receptor abundances, which monovalent ligands, even with engineered affinities, are unable to do.

### Non-Overlapping Ligand Targeting Drives a Limited Selectivity Enhancement of Mixtures

While most therapies rely on the action of a single molecular species, mixtures may enhance selectivity through combinations of actions.^20^ Furthermore, some biologics inevitably act as mixtures of species through heterogeneity in glycosylation and the presence of endogenous ligands.^21^ Therefore, it is important to understand how mixtures of complexes influence overall response.

To evaluate the contribution of mixtures, we evaluated model-predicted binding while varying the composition between two distinct monovalent ligands, each exhibiting preferential binding to either *R*_1_ or *R*_2_(Fig. 4a). The trend that arises is very similar to an additive combination of the single ligand cases. This pattern highlights a key limitation of using mixtures for selectivity: selectivity between two populations varies monotonically with the composition, so any mixture combination is no better than using the more specific ligand entirely (Fig. 4b).

**Figure 4:**
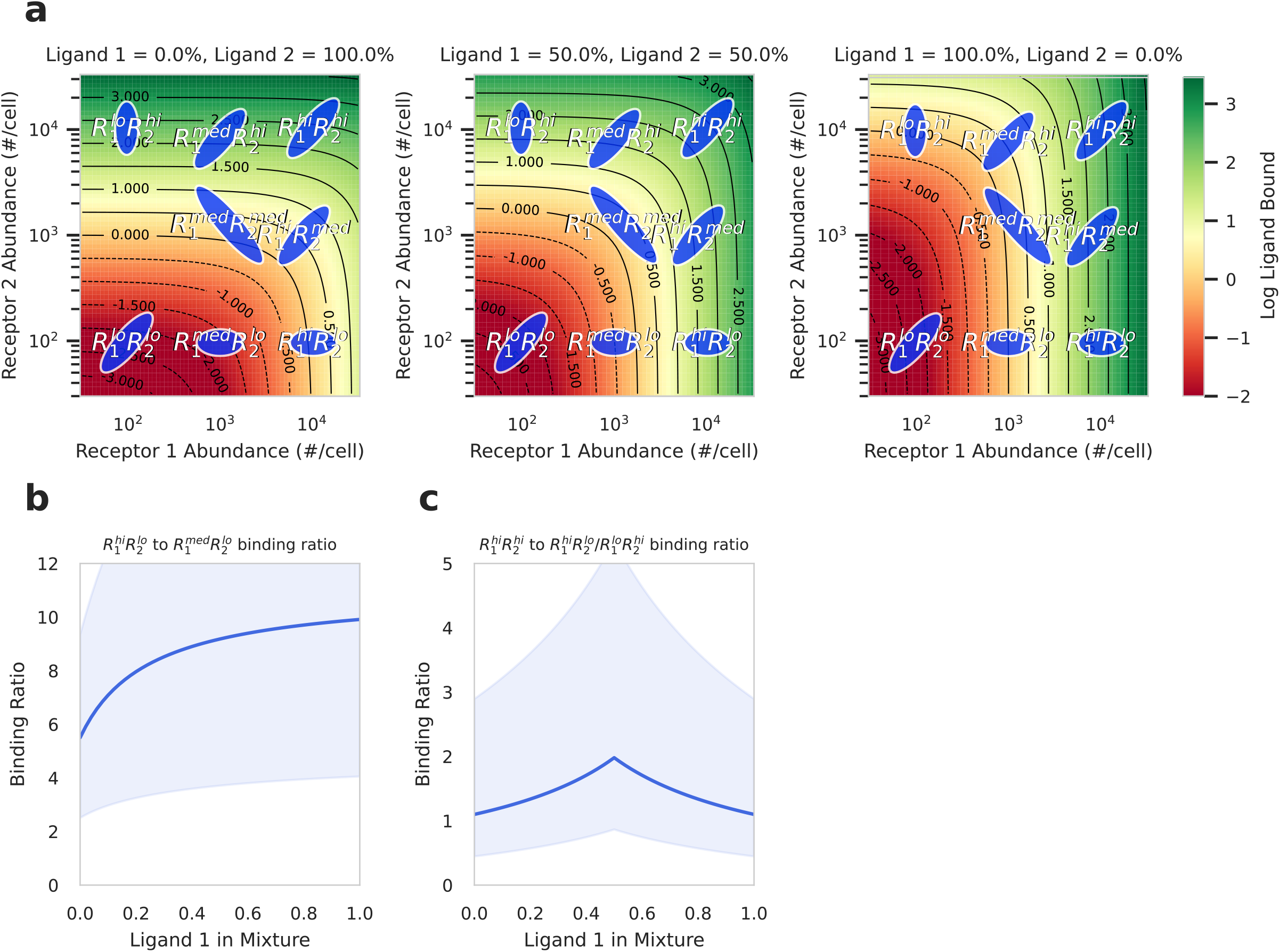
Ligand mixtures with non-overlapping responses can enhance selectivity. a) Heat/contour maps of multivalent ligands bound to cell populations given their expression profiles of two receptors. A mixture of monovalent ligands is used, with ligand 1 binding to receptor 1 and 2 with dissociation constants of 1 *μ*M and 10 *μ*M respectively, and ligand 2 binding to receptors 1 and 2 with dissociation constants of 10 *μ*M and 1 *μ*M respectively. Ligand concentration *L*_0_ = 1 nM; crosslinking constant 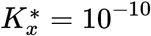. b,c) Ratio of ligand bound to cell populations exposed to monovalent mixtures of ligand 1 and 2. The ratio of the target population to the single off target population with the greatest ligand bound is plotted. The shaded areas indicate the variance caused by intrapopulation heterogeneity, to 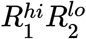 to 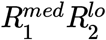, 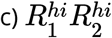 to 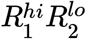 and 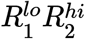.

While mixture engineering fails to enhance binding selectivity between two cell populations, it is potentially beneficial when considering two or more off-target cell populations. More specifically, when a target population expresses two target receptors, but off-target populations express each receptor individually in high amounts, drug mixtures can offer enhanced selectivity. For example, when maximizing targeting to 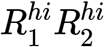 over 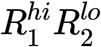 and 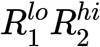, which individually express high levels of the receptors found on the 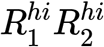, a uniform mixture of ligands with high affinity for receptor 1 and 2 provides a modest improvement in targeting selectivity (Fig. 4c). However, even in these cases, the magnitude of selectivity enhancement is modest. Finally, although we only consider the amount of binding, ligands can have non-overlapping signaling effects even with identical amounts of binding. In these cases, the effect of combinations can be distinct from either individual ligand.^17,22^

### Heterovalent Bispecific Ligands Exhibit Unique Charateristics When Activated Fully Bound

Constructing multispecific drugs has become a promising new strategy for finer target cell specificity with the advancement of engineering techniques.^23^ However, the number of possible configurations of multispecific drugs is combinatorially large and impossible to enumerate. Here, we use heterovalent bispecific ligands as examples to explore the unique benefit of multispecificity distinct from any strategy analyzed before. Here, we compare bispecificity with a 50%-50% mixture of two monovalent ligands, and a 50%-50% mixture of two different homogeneous bivalent ligands (Fig. 5i). These two strategies both have some similarities to bispecific therapeutics. A bispecific ligand contains two different ligand monomers with equal proportion, similar to a 50%-50% monovalent mixture. By comparing bispecific with a 50%-50% monovalent mixture, we can elucidate the extra benefit of tethering these two monomers into one complex. A bispecific ligand is also by nature bivalent, and by comparing it with homogeneous bivalent drugs, we can understand how having two different subunits in the same complex modify the behavior of a drug. Together, we sought to identify the unique advantages afforded by heterovalent ligands.

**Figure 5:**
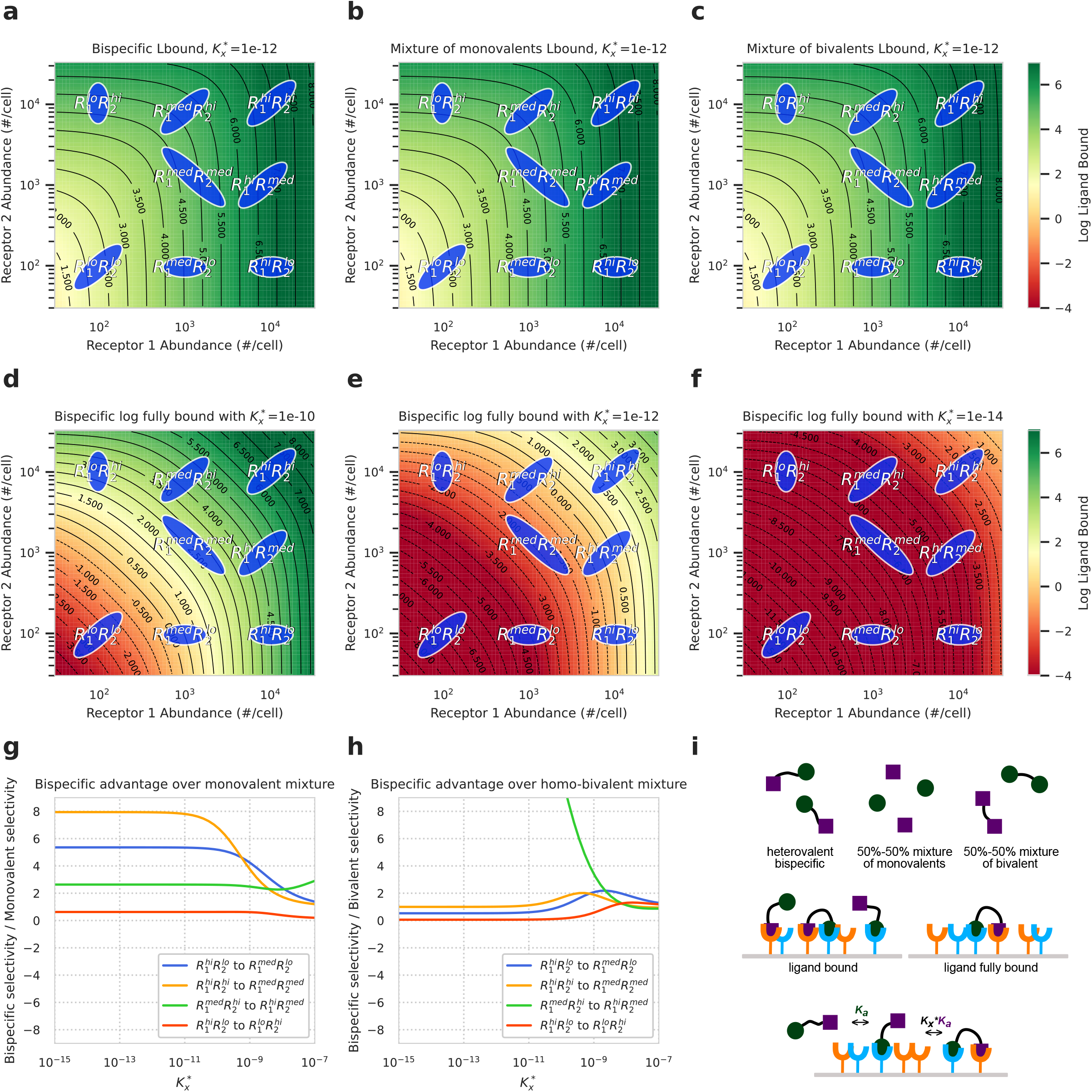
Bispecific ligands exhibit unique effects when they can only be activated with both subunits bind. a-c) Ordinary ligand bound are dominated by the initial binding of ligands, so it doesn’t provide unique advantages to a) bispecific ligand comparing with b) a 50%-50% mixture of monovalent ligands and c) a 50%-50% mixture of bivalent ligands. Ligand concentration *L*_0_ = 10nM; binding affinities *K*_*d*11_ = 100nM, *K*_*d*22_ = 1*μ*M, *K*_*d*12_ = *K*_*d*21_ = 10*μ*M. d-f) The amount of fully bound bispecific ligands depends on the tendency of multimerization, capsuled by 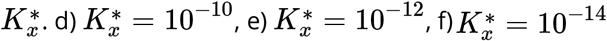. g,h) Comparing bispecific selectivities with mixture selectivities, varies with 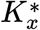, the crosslinking constant. When the ratios are larger, bispecific ligands bind to target populations more specifically. g) bispecific selectivities divided by monovalent 50%-50% mixture selectivities, h) bispecific selectivities divided by a 50%-50% homo-bivalent mixture selectivities. i) Cartoons of bispecific ligands and ligand fully bound binding model.

We first applied the binding model to predict the amount of ligand bound in bispecific drugs (Fig. 5a), a 50%-50% mixture of two monomers (Fig. 5b), and a 50%-50% mixture of two different homogeneous bivalents (Fig. 5c), with the same set of parameters in ligand concentration and affinities. Surprisingly, the patterns of ligand binding in these three cases are almost exactly the same, and bispecificity appeared to offer no unique properties. However, many bispecifics only impart their therapeutic action when both of their subunits bind to a target population. For example, in the design of bispecific antibodies, it is common to require both subunits bound to be effective.^24,25^ When the first antigen-binding region of a bispecific antibody docks at an antigen, it will only make a loose connection between the target and the antibody, unable to induce a strong immune response, and only when both antigen-binding regions bind to their target will this bispecific antibody be fully effective. We investigated whether bispecific ligands that have to be activated with both of their subunits binding to a target display any special cell population selectivity characteristics.

To scrutinize the case for bispecific ligands where the target binding of both subunits is required, we extended our model to calculate the amount of ligand fully bound (Fig. 5i). For the heterovalent bispecific case, ligand fully bound will only account for the amount of ligand with both of their ligand monomers bound to a target. With the same set of parameters, the predictions made for bispecific fully bound show a very distinct pattern from general ligand bound (Fig. 5e). The contour plot of fully bound bispecific ligands has more convex contour lines. For example, 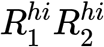 has about the same level of general ligand bound as 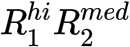 (Fig. 5a), but it has significantly more ligand fully bound than 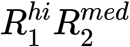 (Fig. 5e). This convexity of contour lines indicates that for bispecific complexes, double-positive cells bind more ligands fully than cells only strongly expressing one type of receptors. This should be obvious since these two subunits of the bispecific complex prefer different receptors. The results indicate that bispecific ligands will only exhibit special characteristics when it requires both receptor binding subunits to bind a receptor.

The specific amount of fully bound ligands is dependent on the ligand’s propensity for crosslinking captured by the constant 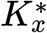 (Fig. 5i). Typically in biochemistry, when there is less steric hindrance among the subunits of a multivalent drug molecule (e.g. longer tether and smaller subunit size), or when there is local receptor clustering on the target cell, secondary binding will be easier to achieve, corresponding to larger 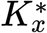.^26^ We plotted the pattern of bispecific fully bound with 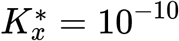,10^−12^, and 10^−14^ (Fig.5d-f). In general, when 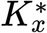 is larger, the ligands are more capable of multimerization, and there will be more fully bound units. To demonstrate how this characteristic of fully bound bispecific ligand imparts cell population specificity, we compared the selectivities between some chosen target-to-off-target cell population pairs under bispecific drug versus another drug given a range of 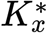 (Fig. 5g,h). These plotted numbers are the selectivities imparted by a bispecific drug divided by the selectivities from a drug mixture of either monovalent (Fig. 5g) or homogeneous bivalent (Fig. 5h). When these quotients are larger, it implies that fully bound bispecific ligand has greater advantages than its counterpart to impart selective binding for the chosen target cell populations. Figure 5g compares the selectivities under bispecific ligands versus a 50%-50% mixture of monovalent ligands. The results show that fully bound bispecific can grant better selective binding when 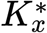 is small enough. This is logical, since when 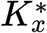 is small and crosslinking is rarer, most ligands will bind monovalently, and fully bound bispecific will especially favor the cell populations with higher receptor expression. However, when we compare fully bound bispecific to fully bound homogeneous bivalent mixtures (Fig. 5h), the advantage of bispecific drugs is not obvious except for 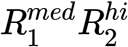 to 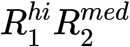 selectivity. Given that we only account for fully bound ligand for both therapeutics, the effect demonstrated in Figure 5g no longer holds. Together, we showed that bispecific ligands only exhibit unique advantages in inducing selective binding when they are only effective when both of their subunits bind and crosslinking is more difficult.

### Combining Strategies For Superior Selectivity

Each strategy described above provided selectivity benefits in distinct situations, suggesting that they might synergistically improve selectivity when combined. We explored this potential synergy using optimization to determine the ligand specifications which provided optimal selectivity for our theoretical cell populations. More specifically, we optimized the selectivity of a ligand for a particular population, while considering all other populations to be off-target. Our optimization allowed ligand characteristics to vary within biologically plausible bounds; which characteristics we allowed to vary defined our examination of the efficacy of our various ligand engineering strategies. We examined optimizing affinity alone, mixture along with affinity, and valency along with affinity, and finally combined all three strategies (Fig. 6).

**Figure 6:**
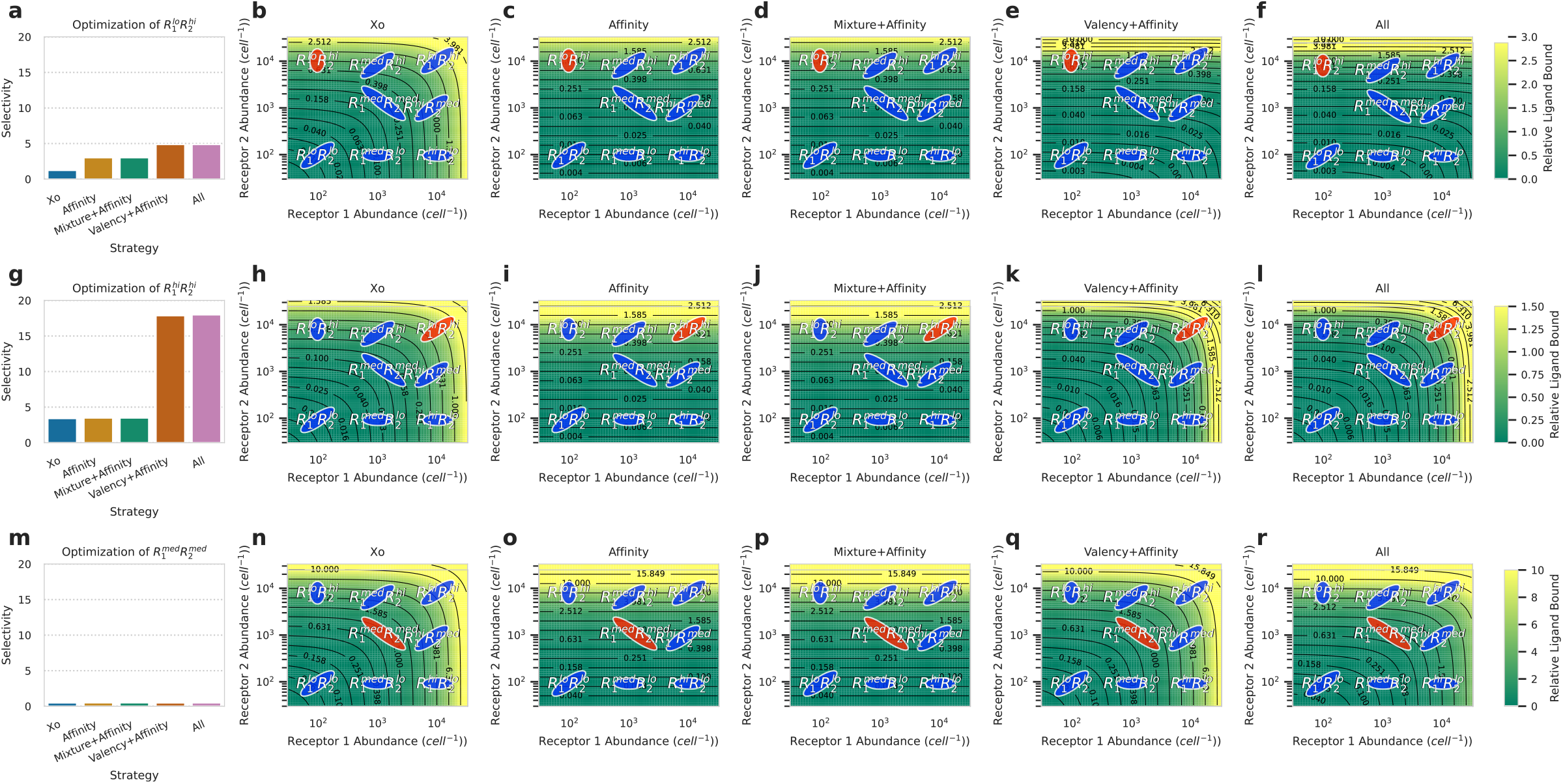
Combinations of strategies provide superior selectivity. a,g,m) Optimal selectivity levels (ligand bound on target population divided by average ligand bound by all other populations) achieved using various ligand engineering techniques. Ligand concentration *L*_0_ = 1nM. Xo ligands are monovalent ligands with affinities of 1 *μ*M for both receptor 1 and 2. The dissociation constant was allowed to vary between 10 mM and 0.1 nM for both receptors using the “affinity” approach. Valency was allowed to vary from 1 to 16 for the “valency” approach in addition to affinities varying. Mixtures were assumed to be composed of two monovalent ligands, and affinities were allowed to vary in the “mixture” approach. The combined “all” approach allowed all of these quantities to vary simultaneously. The crosslinking constant 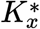 was allowed to vary between 10^−15^ and 10^−9^ for all approaches. b-f,h-l,n-f) Heatmap of magnitude of ligand bound for ligand with optimized characteristics according to various ligand engineering strategies. Target population is shown in red. a-f) Pertains to optimal targeting of 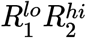, g-l) pertains to optimal targeting of 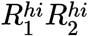, and m-r) pertains to optimal targeting of 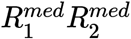.

Optimizing a ligand for selectivity to 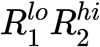 highlights a situation in which affinity imparts greater specificity, and optimal selectivity is achieved by combining affinity and valency modulations (Fig. 6a-f). Here selectivity is optimized by ligands with selective binding to receptor 2, and those with valencies allowing for selective binding to cells with higher abundances of receptor expression. One case contradictory to this trend is shown during the optimization for selectivity towards the 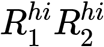 (Fig. 6g-l). While affinity engineering is unable to impart some small contributions to enhanced selectivity, significant improvement is only achieved when utilizing valency modulation techniques. A more difficult design problem is featured in the optimization of 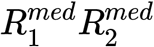 (Fig. 4m-r). Since it lies in the midst of the other populations in receptor expression space, any modulation of affinity, valency or combining it with mixture-based strategies seems ineffective. It should be noted that in all described cases, engineering the composition of a mixture of ligands is generaly ineffective for imparting selectivity when the ligand’s design specifications are flexible, and is likely only efficacious when using ligands with static properties and considering multiple off-target populations.

Our results highlight that both in singular and combined strategies for therapeutic manipulation, the target and off-target populations dictate the optimal approach. It is also clear that combined approaches do offer synergies which can be harnessed, but that those are only emergent in particular therapeutic situations.^25^

### Using Binding Competition to Invert Receptor Targeting

While the strategies above provided selectivity in many cases, we recognized that they are all limited to a positive relationship between receptor abundance and binding. Therefore, we wondered if binding competition with a receptor antagonist, or “dead ligand”, could invert this relationship.

To investigate the effect of ligand competition with an antagonist, we modeled mixtures of ligands, but only quantified the amount of binding for the active ligand. Here we chose to only consider a monovalent agonist and tetravalent antagonist (Fig. 7). We found that combinations of monovalent agonistic ligands and multivalent antagonistic ligands were able to uniquely target populations expressing small or intermediate amounts of receptors, which is demonstrated when comparing ligand binding ratios between 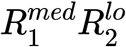 to 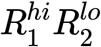 (Fig. 7a). Here, a nearly tenfold increase in selectivity can be granted to monovalent agonists when combined with a tetravalent antagonist. In this case, there are greater quantities of agonist bound to 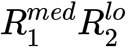 than 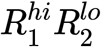 (Fig. 7b). This is striking, as 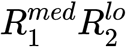 expresses either as many or fewer abundances of receptors one and two when compared to 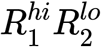. This phenomenon, which could not be achieved without multivalent antagonists, occurs due to the preferential binding of multivalent antagonists to populations expressing higher abundances of receptors (Fig. 7c, 3e). Thus, in cases where previously discussed ligand engineering strategies and approaches fail to achieve selective binding to cells expressing smaller or similar amounts of receptors to off-target populations, combinations of agonistic and antagonistic ligands may provide unique benefits.

**Figure 7:**
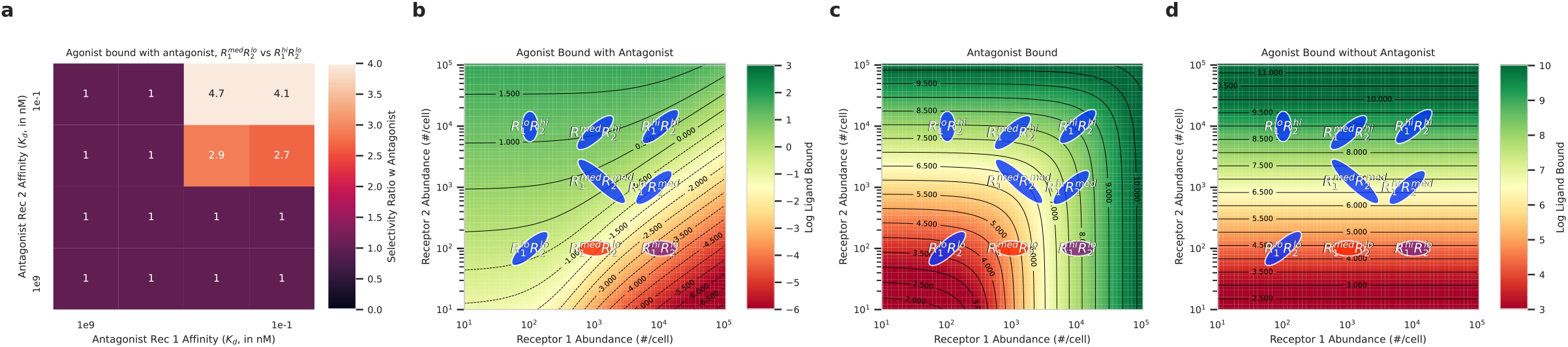
Mixtures of receptor agonists and antagonists allow for unique population targeting activity. Ligand concentration *L*_0_ = 1nM. a) Selectivity for 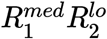 against 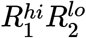 when exposed to a tetravalent “dead ligand” antagonist with varying affinities for receptors 1 and 2, and a monovalent therapeutic receptor agonists with affinities optimized for selectivity. Only amount of agonist bound is considered in determination of optimal selectivity. b-d) Heatmap of agonist (b), and antagonist (c) ligand bound for antagonist and agonist ligand combination shown to impart greatest selectivity improvement in (a). d) Heatmap of agonist bound in b,c when no antagonist is present.

**Figure 8:**
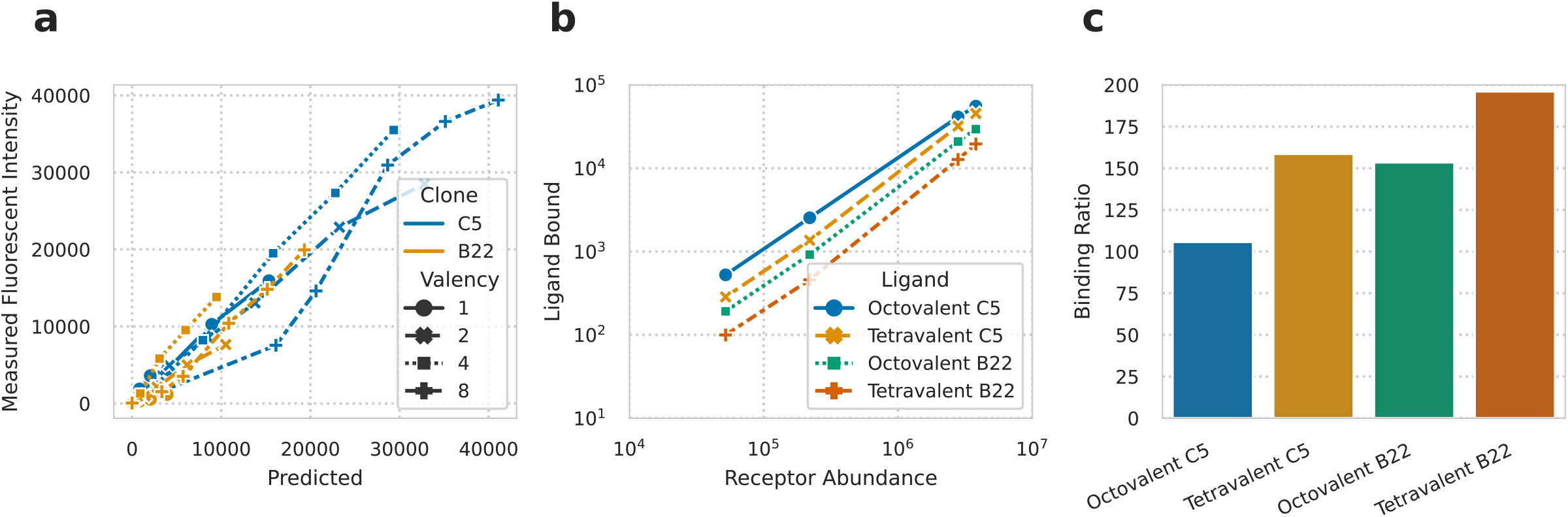
Our model is able to recapitulate experimental multivalent binding activity. a) Experimental vs. predicted fluorescent units for MCF-7 cells expressing 3.8 × 10^6^ fibronectin receptors per cell bound to nanorings expressing one, two, four or eight fibronectin binding domains at concentrations of 0.16– 500 nM. C5 and B22 are high and low affinity fibronectin binding domains respectively. Using nonlinear least-squares regression, the crosslinking coefficient was found to be 1.11 × 10^−12^; the dissociation constants for C5 and B22 were found to be 1.73 *μ*M and 3.24 *μ*M respectively, and ligand to fluorescent conversion factor for C5 and B22 were found to be 5.1 × 10^−2^ and 3.2 × 10^−2^ respectively. b) Predicted fluorescent values for MDA cells expressing 5.2 × 10^4^, SK cells expressing 2.2 × 10^5^, LNCaP cells expressing 2.8 × 10^6^, and MCF-7 cells expressing 3.8 × 10^6^ receptors per cell as described in Csizmar et al. ^18^. Fluorescence was predicted for interactions with octa- and tetravalent ligands expressing either C5 or B22 fibronecting binding domains. c) Predicted binding ratios of MCF-7 cells to MDA cells when bound with octa- and tetravalent ligands expressing either C5 or B22 fibronecting binding domains.

## Discussion

Here, we developed and employed a multivalent, multi-ligand, multi-receptor binding model and used it to explore the effectiveness of various ligand engineering strategies for population-selective binding (Fig. 1). Using a representative set of theoretical cell populations defined by their distinct expression of two receptors, we examined the efficacy of several potential ligand engineering strategies, including changes to affinity, ligand valency, mixtures of species, multi-specificity, antagonist competition, and these in combination.

Each strategy’s contribution can be summarized by general patterns. We found that affinity changes were most effective when the target and off-target populations expressed distinct combinations of receptors (Fig. 2). Binding selectivity was enhanced by increasing the affinity of the ligand for those receptors that are more abundantly expressed on the target cells and decreasing its affinity for those that are not. Selectivity depended upon large differences in receptor expression levels between on- and off-target populations. When target and off-target populations expressed the same pattern of receptors and only differed in receptor abundances, modulations in valency, but not affinity, were effective (Fig. 3). A key determinant of valency’s effectiveness was the secondary binding and unbinding rate which is dependent on both the kinetics of the receptor-ligand interaction and receptor abundance. Ligand mixtures were mostly ineffective for imparting binding selectivity, and only had modest benefits when considering two or more off-target populations (Fig. 4). Heterovalent bispecific ligands only showed unique advantages over mixtures of monovalent ligands or bivalent ligands when solely considering fully bound ligand (Fig. 5). These ligands exhibit preferential binding to target populations that have high expression of both receptors over those with high expression of only a single receptor, with the propensity for secondary binding acting as a key determinant for selectivity.

We found that, while a single ligand engineering strategy dominated in its contributions to cell type selectivity, synergies between these strategies existed in some cases to derive even greater specificity (Fig. 6). Finally, we found that combinations of monovalent therapeutic ligands with multivalent antagonistic ligands allow for the unique selection of target populations expressing relatively fewer receptors than off-target populations (Fig. 7).

While our multivalent binding model identified strategies for selective targeting in many cases, it also identified situations for which selective binding is challenging. For example, selectively targeting populations based on their absence of receptor expression remains challenging. While we computationally show the potential of using multivalent antagonists with monovalent agonists to selectively target such populations, implementing this may be challenging in practice. In cases where a target population expresses fewer receptors of any kind than an off-target population, our analysis suggests that targeting other receptors should be considered. However, in cases where target populations express more of any type of receptor than an off-target population, we show that one or more of our formulated ligand engineering strategies can be employed to improve binding selectivity. While we expect the same patterns to apply with greater than two receptors, certain emergent behaviors may exist with tri-specific and more complex ligand binding.

A few of the strategies that we explored have been utilized in existing engineered therapies. For example, affinity changes to the cytokine IL-2 have been used to bias its effects towards either effector or regulatory immune populations.^27,28^ Varying the valency of tumor-targeted antibodies leads to selective cell clearance based upon the levels of expressed antigen.^19^ Manipulating of the affinities of the fibronectin domains on octovalent nanorings was shown to enhance the selectivity of binding to cancerous cells displaying relatively higher densities of fibronectin receptors compared to native tissue.^18^ The tendency of low-affinity, multivalent interactions to target cells expressing high receptor abundances was also described in a study describing the selectivity of multivalent antibody binding to tumor cells bound by a bispecific therapeutic ligand.^13^ These examples lend support to the accuracy and translational value of our model. At the same time, recognizing these previously described ligand engineering approaches as separable strategies provides clearer guidance for future engineering.

Some of the optimization strategies described here have not been exploited in part due to the complexity of real biological applications. First, some strategies may not be biochemically practical. For example, the manipulating ligand affinity requires intricate protein engineering. Potential dynamic changes in the receptor expression profile of a target population also complicate the matter. It is well documented that cancer cells can escape therapeutic targeting by upregulating^29,30^ or downregulating^31^ the expression of certain receptors. In this case, both the current and potential abundance of each receptor must be considered. While this work does not address many of these issues, we propose that using a computational binding model can analyze these strategies quantitatively and collectively from a mechanistic perspective. Even when the absolute mathematically optimal ligand characteristics cannot be achieved biochemically, our analyses provide guidance within the bounds of what is attainable and how to approach the optimum, accounting for implementation feasibility and facilitating the implementation of strategy combinations.

In many therapeutic applications where selective engagement of target cell populations is an important performance metric, such as the treatment of cancer, cellular therapies are becoming increasingly popular.^32^ Human engineered chimeric antigen receptor (CAR) T cells have enhanced the potential to selectively recognize and attack malignant tissues.^33^ These technologies bypass ligand-receptor binding restrictions by allowing recognition in signaling response. However, we have shown here that selectivity can often be attained with relatively simple therapeutic ligands. This study lays the framework for how ligand engineering can be directed using computational modeling. It should be noted that the application of this logic is reliant on knowledge of the target and off-target cell population receptor expression levels. Future application of the ligand binding logic described in this study could be guided using high-throughput single-cell profiling techniques, such as RNA-seq or high-parameter flow cytometry. A computational tool that could directly translate such datasets into ligand design criteria, selecting among potential receptor targets, may represent a potential avenue for the translation of our analyses into a more broadly applicable ligand engineering tool.

## Materials and Methods

### Data and Software Availability

All analysis was implemented in Python v3.8, and can be found at https://github.com/meyer-lab/cell-selective-ligands.

### Generalized multi-ligand, multi-receptor multivalent binding model

To model multivalent ligand complexes, we extended our previous binding model to the multi-ligand case.^17^ We define *N*_*L*_ as the number of distinct monomer ligands, *N*_*R*_ the number of distinct receptors, and the association constant of monovalent binding between ligand *i* and receptor *j* as *K*_*a,ij*_. Multivalent binding interactions after the initial interaction have an association constant of 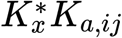 proportional to their corresponding monovalent affinity. The concentration of complexes is *L*_0_, and the complexes consist of random ligand monomer assortments according to their relative proportion. The number of ligand complexes in the solution is usually much greater than that of the receptors, so we assume binding does not deplete the ligand concentration. The proportion of ligand *I* in all monomers is *C*_*i*_. By this setup, we know 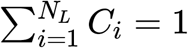. *R*_tot,*i*_ is the total number of receptor *I* expressed on the cell surface, and *R*_eq,*i*_ the number of unbound receptors *i* on a cell at the equilibrium state during the ligand complex-receptor interaction.

The binding configuration at the equilibrium state between an individual complex and a cell expressing various receptors can be described as a vector **q** = (*q*_1,0_, *q*_1,1_,…, *q*_1,*NR*_, *q*_2,0_,…, *q*_2,*NR*_, *q*_3,0_,…, *qN*_*L*_,*N*_*R*_) of length *N*_*L*_(*N*_*R*_ + 1), where *q*_*i,j*_ is the number of ligand *i* bound to receptor *j*, and *q*_*i*,0_ is the number of unbound ligand *i* on that complex in this configuration. The sum of elements in **q** is equal to *f*, the effective avidity. For all *i* in {1, 2,…, *N*_*L*_}, let 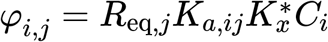 when *j* is in {1, 2,…, *N*_*R*_}, and *φ*_*i*,0_ = *C*_*i*_. The relative amount of complexes in the configuration described by **q** at equilibrium is

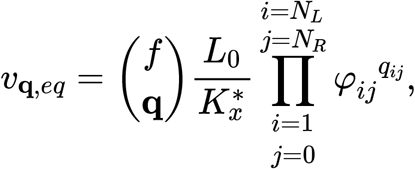

with 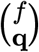 being the multinomial coefficient. Then the total relative amount of bound receptor type *n* at equilibrium is

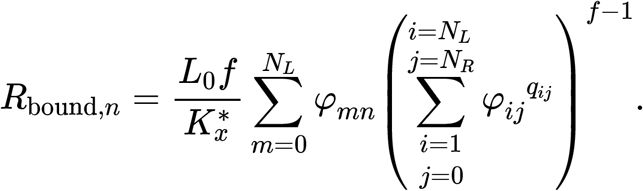

By conservation of mass, we know that *R*_tot,*n*_ = *R*_eq,*n*_ + *R*_bound,*n*_ for each receptor type *n*, while *R*_bound,*n*_ is a function of *R*_eq,*n*_. Therefore, each *R*_eq,*n*_ can be solved numerically using *R*_tot,*n*_. Similarly, the total relative amount of complexes bound to at least one receptor on the cell is

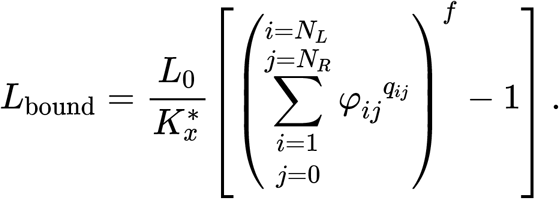

### Generalized multivalent binding model with defined complexes

When complexes are engineered and ligands are not randomly sorted into multivalent complexes, such as with the Fabs of bispecific antibodies, the proportions of each kind of complex become exogenous variables and are no longer decided by the monomer composition *C*_*i*_’s. The monomer composition of a ligand complex can be represented by a vector 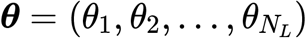, where each *θ*_*i*_ is the number of monomer ligand type *i* on that complex. Let *C*_***θ***_ be the proportion of the ***θ*** complexes in all ligand complexes, and Θ be the set of all possible ***θ***’s. We have ∑_***θ***∈Θ_ *C*_***θ***_ = 1.

The binding between a ligand complex and a cell expressing several types of receptors can still be represented by a series of *q*_*ij*_. The relationship between *q*_*ij*_’s and *θ*_*i*_ is given by *θ*_*i*_ = *q*_*i*0_ + *q*_*i*1_+… +*q*_*iNR*_. Let the vector **q**_*i*_ = (*q*_*i*0_, *q*_*i*1_,…, *q*_*iNR*_), and the corresponding ***θ*** of a binding configuration **q** be ***θ***(**q**). For all *i* in {1, 2,…, *N*_*L*_}, we define 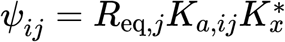 where *j* = {1, 2,…, *N*_*R*_} and *ψ*_*i*0_ = 1. The relative amount of complexes bound to a cell with configuration **q** at equilibrium is

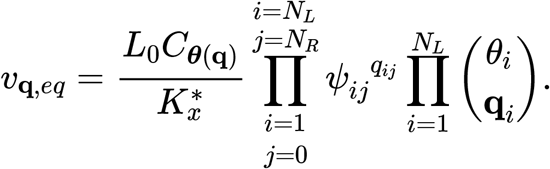

Then we can calculate the relative amount of bound receptor *n* as

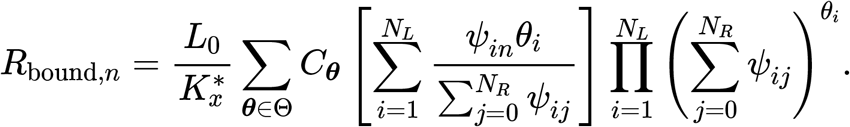

By *R*_tot,*n*_ = *R*_eq,*n*_ + *R*_bound,*n*_, we can solve *R*_eq,*n*_ numerically for each type of receptor. The total relative amount of ligand binding at equilibrium is

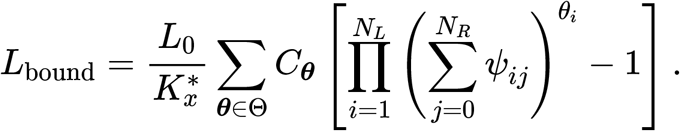

### Mathematical optimization

We used the SciPy function scipy.optimize.minimize to combine several strategies and achieve optimal selectivity.^34^ Unless specified otherwise, the initial values for optimization were 10^−12^ for crosslinking coefficient 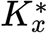, 1 for valency *f*, 100% ligand 1 for mixture composition, and 1 *μ*M for the affinity dissociation constants. The boundaries were 10^−15^ to 10^−9^ for 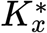 for *f*, 0–100% ligand 1 for mixture composition, and 10 mM to 0.1 nM for the affinity dissociation constants.

### Sigma point filter

To consider the intrapopulation variance of a cell population in the optimization, we implemented the sigma point filters,^16^ a computationally efficient method to approximate the variance proprogated through an ordinary differential equation-based model.

### Reimplementation of Csizmar et al

To validate our model, we recapitulated multivalent binding data from Csizmar et al. using our multivalent binding model (Fig. S1).^18^ Here, fluorescently labeled nanorings displaying 1, 2, 4, and 8 fibronectin clones, which bind to epithelial cell adhesion molecule (EpCAM) antigens, were fabricated. Binding activity of nanorings displaying both high (C5) and low (B22) affinity EpCAM binding domains was measured. Binding to an EpCAM^high^ (3.3 × 10^6^ antigens/cell) population was measured using flow cytometry. We used nonlinear least squares optimization, as described above, to fit our multivalent binding model to the binding data, using a unique scaling factor for each fibronectin clone to convert between measured fluorescent intensity and magnitude of ligand bound. We allowed affinity of the fibronectin clones to vary during optimization.

## Supplementary Figures

## Acknowledgements

This work was supported by NIH U01-AI148119 to A.S.M.

## Author contributions statement

A.S.M. concieved of the work. All authors implemented the analysis and wrote the paper.

